# Comparative transcriptome analysis of the ovary and testis of the lacustrine goby (*Gobiopterus lacustris*)

**DOI:** 10.1101/781104

**Authors:** Zhongdian Dong, Chengqin Huang, Hairui Zhang, Shunkai Huang, Ning Zhang, Changxu Tian, Zhongduo Wang, Yusong Guo

## Abstract

Lacustrine goby (*Gobiopterus lacustris*) belongs to a genus of gobies that are small in size and endemic to freshwater, brackish waters or coastal environments around the Indian and Pacific oceans. To date, there are no genomic or transcriptomic studies on *G. lacustris*. Here, we constructed gonadal transcriptomes of *G. lacustris* for the first time and identified genes that may be involved in gonadal development and reproduction. In total, 60,657,644 and 52,016,136 clean reads were obtained from ovary and testis, respectively, using Illumina sequencing. Reads were assembled into 62,573 unigenes with N50 value of 3,082 bp and a mean length of 1,869 bp. A total of 47,891 (76.53%) unigenes were annotated in at least one of the seven databases that were used in this study. In addition, 38,550 SSRs (simple sequence repeat, microsatellite) were identified from 20,517 SSR containing sequences. Gene expression patterns in the testis and ovary were compared, and 10,954 DEGs (differentially expressed genes) were identified. Of these genes, 2,383 were up-regulated in the testis and 8,571 were up-regulated in the ovary. RT-qPCR analysis of 14 selected genes showed patterns consistent with the transcriptome results. Numerous DEGs involved in gonadal development and gametogenesis were identified, including *foxl2*, *dmrt1*, *cyp19a1a*, *inha*, *inhb*, *sycp2*, *zglp1*, *tdrp, zps* and *esra*. Using GO and KEGG enrichment analyses, pathways involving regulation of gonadal development and gametogenesis also identified. This work represents the first gonadal transcriptomic analysis of *G. lacustris* and provides a valuable dataset for future research on the genes involved in reproduction of *G. lacustris*.

## 1. Introduction

Sex differentiation of fish is determined by both genetic and environmental factors. Only a few fish species have master sex-determination genes [1, 2]. These genes include *dmy* (Y-specific DM-domain) from Japanese medaka (*Oryzias latipes*) [3], *sdY* (sexually dimorphic on the Y chromosome) from rainbow trout (*Oncorhynchus mykiss*) [4], *amhr2* (*anti*-Mullerian hormone receptor type II) from fugu (*Takifugu rubripes*), *amhy* from patagonian pejerrey (*Odontesthes hatcheri*) [5], *gsdf* (gonadal soma-derived growth factor on the Y chromosome) from *Oryzias luzonensis* [6] and *dmrt1* (Doublesex and mab-3 related transcription factor 1) from Chinese tongue sole (*Cynoglossus semilaevis*) [7, 8]. In addition to master sex determining genes, conserved genes such as *cyp19a1a, foxl2*, *fox3*, *igf3, sox9*, and *sf1* are involved in sex determination of fish [9–12]. Biological pathways, such as the TGF-beta signaling pathway and estrogen signaling pathway, also pay essential roles in sex determination and gonadal development [1, 13, 14]. To better understand the regulatory mechanisms underlying sex determination in fish, it is essential to explore genes involved in this process at a broader phylogenetic context.

Transcriptomes allow for general characterization of genes that are expressed in specific tissues or under specific conditions and have been successful in identifying sex-related genes and associated regulatory mechanisms in fish [9, 13]. Genes involved in gonad development and gametogenesis have been identified using transcriptomes in many aquatic species, including the channel catfish [15], olive flounder [9, 16], Japanese scallop [16], spotted knifejaw [13], Chinese giant salamander (*Andrias davidianus*) [17] and silver sillago (*Sillago sihama*) [14] .

Lacustrine goby (*Gobiopterus lacustris*) is a bony fish that is smaller than zebrafish. The body of *G. lacustris* is translucent or transparent, and this species has the potential to be a model organism for pigmentation. We recently found that *G. lacustris* is endemic to Luzon, Philippines, and is abundant in the mangroves of Leizhou Peninsula, Guangdong Province, China [18, 19]. In this study, we generated the gonadal transcriptome of *G. lacustris* and identified genes that may be involved in gonadal development and reproduction. The transcriptome will provide information for further research on the regulatory mechanisms of sex determination and differentiation in *G. lacustris*.

## 2. Material and methods

### 2.1. Identification of gonadal development, RNA isolation, cDNA Library preparation and sequencing

The *G. lacustris* (body length 18.8 ± 0.5 mm; body weight 0.11 ± 0.03 g) were collected from Zhanjiang Mangrove National Nature Reserve, Guangdong Province, China, and were domesticated for three months in the laboratory at 15 ‰ salinity. In order to define the developmental state of the gonads, we prepared gonad tissue sections. Three female and three male mature *G. lacustris* were collected and fixed in Bouin’s liquid for 18 hours and dehydrated in an ethanol series, cleared in xylene, and embedded in paraffin. Sections were cut at 8 μm thickness and stained with hematoxylin- eosin. Three sexually mature females and males each were selected and placed on ice to dissect the ovary and testis for RNA extraction. All experimental protocols were approved by the Animal Research and Ethics Committee of Guangdong Ocean University (NIH Pub. No. 85-23, revised 1996). The total RNA of mature gonads was extracted using Trizol reagent (Invitrogen, Carlsbad, USA) following the manufacturer’s protocol. Genomic DNA was removed using DNase I (TaKaRa, Dalian, China). RNA degradation and contamination were monitored using 1% agarose gels, RNA integrity was assessed using the RNA Nano 6000 Assay Kit for the Bioanalyzer 2100 system (Agilent Technologies, CA, USA) and RNA concentration was measured using Qubit® RNA Assay Kit in Qubit® 2.0 Flurometer (Life Technologies, CA, USA).

For each sex, 1 μg total RNA was taken from each of the three samples and mixed to create a pooled sample. A total amount of 1.5 μg total RNA per sample was used as input material for transcriptomic analysis. Sequencing library preparation and sequencing were completed by Novogene (Tianjin, China) following methods from Nan et al. (2018). All raw data have been submitted to the CNGB Nucleotide Sequence Archive (CNSA) under the accession number CNP0000359.

### 2.2. Quality control and assembly

Raw data (raw reads) in fastq format were processed through in-house perl scripts. Clean data (clean reads) were obtained by removing reads containing adapter or ploy-N (N indicates that the base information cannot be determined) sequences, as well as low quality reads. The Q20, Q30 and GC content of the clean data were calculated. All the downstream analyses were conducted on the clean data to ensure high quality data. Transcriptome assembly was accomplished based on the clean data using Trinity-v2.5.1 (Trinityrna-seq_r20140413p1) with min_kmer_cov set to 2 by default and all other parameters set default [20].

### 2.3. Functional annotation

The following public databases were used to annotate homologous genes using the BLAST program with E-value cut-off of 1E^−5^: NR (NCBI non-redundant protein databases, http://www.ncbi.nlm.nih.gov), Nt (NCBI non-redundant nucleotide sequences, http://www.ncbi.nlm.nih.gov), Swiss-Prot (A manually annotated and reviewed protein sequence database, http://www.expasy.ch/sprot), KO (KEGG Ortholog database, https://www.kegg.jp/kegg/ko.html), COG/KOG (Clusters of Orthologous Groups of proteins, http://www.ncbi.nlm.nih.gov), GO (Gene Ontology, http://www.geneontology.org) and KEGG (Kyoto Encyclopedia of Genes and Genomes, http://www.genome.jp/kegg). Protein functional annotations were obtained based on alignment results. The results from the Blast search were imported into Blast2GO to assign GO terms [21]. KEGG Automatic Annotation Server (KAAS) was applied to complete KEGG annotation [22]. Pfam (Protein family) annotation was based on hmmscan in the HMMER 3.0 package with 1E^−5^ [23, 24].

### 2.4. Differential gene expression analysis

The transcriptome obtained by splicing Trinity was used as a reference sequence (ref), and the clean reads of each sample were mapped onto the ref. In this process we used RSEM software (bowtie2 default parameter) [25]. Prior to differential gene expression analysis, for each sequenced library, Tthe read counts for each library was adjusted using edgeR 3.0.8 through one scaling normalized factor prior to differential gene expression analysis [26]. Differential expression analysis was performed using the DEGseq (2010) R package. Pvalue was adjusted using q value [27]. q value < 0.005 & |log2(foldchange)| > 1 was set as the threshold for significantly differential expression.

### 2.5. RT-qPCR validation

Fourteen genes were selected based on their transcriptomic expression profile in testis and ovary for validation using RT-qPCR. Primers were designed according to the transcriptome sequences (Table s1). RT-qPCR was performed using Roche LightCycler 96 (Roche, Forrentrasse, Switzerland) with SYBR Premix Ex Taq II (TaKaRa, Dailian, China). The reaction was carried out at a final volume of 15.0 μL, containing 7.5 μL 2 × SYBR Premix Ex Taq II, 0.6 μL of each forward and reverse primer, 1.5 μL cDNA and 4.8 μL ddH_2_O. PCR amplification as follows: 30 s at 95°C for pre-incubation, followed by 40 cycles at 95 °C (5 s), 58 °C (30 s) and 72 °C (30 s). eef1b and hprt1 are commonly used as reference genes in fish [28, 29] and they were stably expressed in the gonadal transcriptome of G. lacustris. So *eef1b* and *hprt1* were used as reference genes to determine relative expression. and three technical repetitions for each sample. Target gene expression was analyzed using the 2^−ΔΔCt^ method and the relative expression level of male gonad used as control sample [30]. Statistical analysis was performed using SPSS 17.0.

### 2.6. Single sequence repeat (SSR) locus identification

The MIcroSAtellite (MISA) (http://pgrc.ipk-gatersleben.de/misa/) was employed to mine for microsatellites in the unigenes. The minimum number of repeat units for mono- nucleotide di-nucleotide, tri- nucleotide, tetra- nucleotide, penta- nucleotide and hexa-nucleotide motifs were set as 10, 6, 5, 5, 5 and 5, respectively. Based on the MISA analysis, Primer 3 was used to design primer pairs in the flanking regions of SSRs for subsequent validation. Separation distance of two SSRs >100 bases were considered separate SSRs.

## 3. Results

### 3.1. Gonadal development, Illumina sequencing and de novo assembly

To better understand the gonadal development of *G. lacustris,* we conducted histological analysis of the male and female gonads (Figure 1). The gonads are symmetrically distributed on both sides of the body cavity. Based on the morphology of the gonads, *G. lacustris* used in this study were determined to be sexually mature.

**Figure 1.**
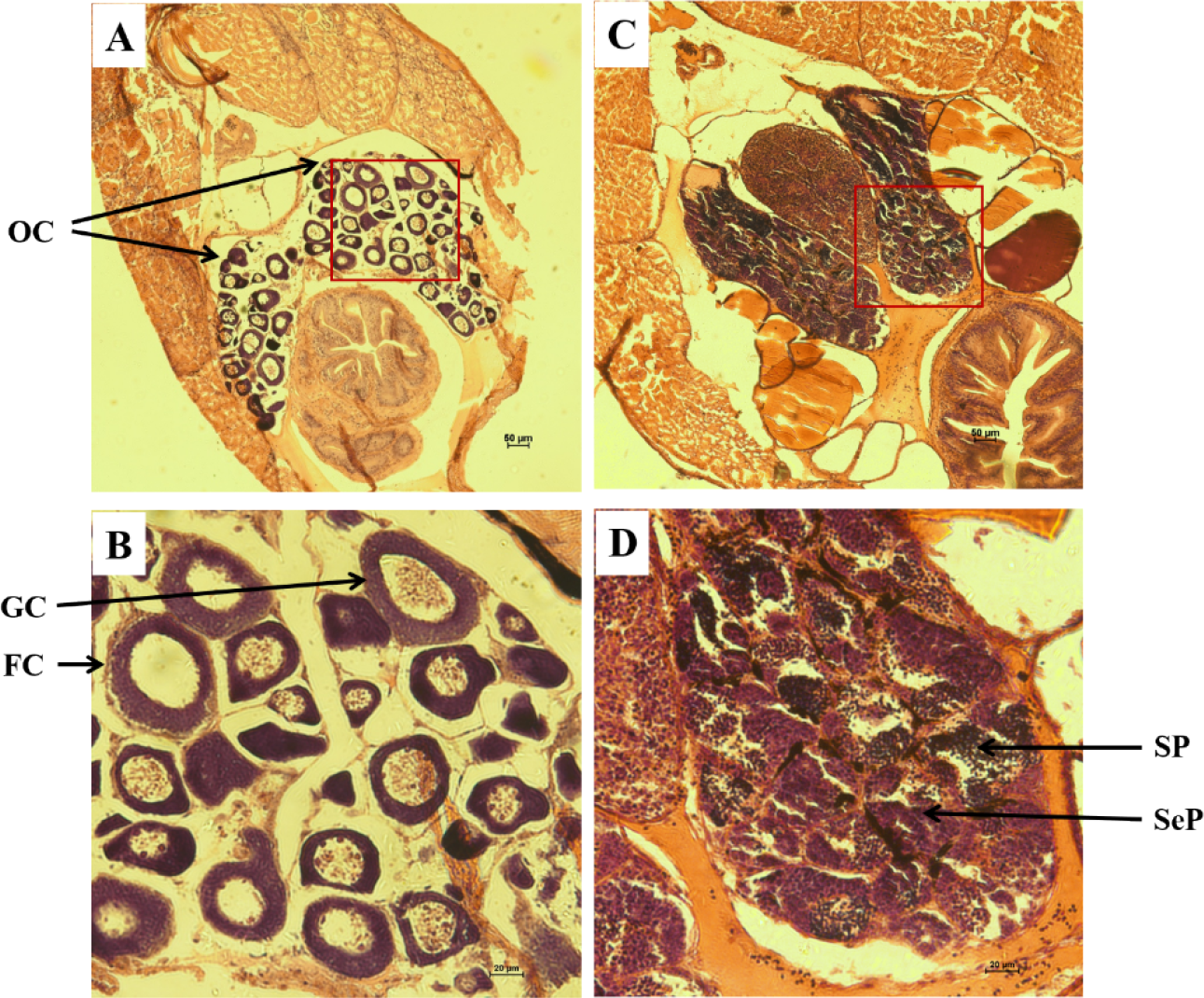
Identification of gonadal development of *G. lacustris*. (A-B) Histology sections of the female gonad. (C-D) Histology sections of the male gonad. Abbreviations are as follows: OC ovarian cavity, GC oocyte, FC follicle cells, Sp spermatid and SeP secondary spermatocyte.

cDNA libraries for the ovary and testis were constructed, and transcriptome sequencing was performed on the Illumina Hiseq platform. A total of 63,391,676 and 54,542,800 raw reads were obtained for the ovary and testis, respectively (Table 1). After performing quality control, 60,657,644 clean reads were retained for the ovary and 52,016,136 clean reads were retained for the testis. The assembled transcriptome had 67,057 transcripts with N50 length of 3,037 bp and 62,573 unigenes with N50 length of 3,082 bp (Table s2). There were 35,081 (56.06% of 62,573) unigenes exceeding 1,000 bp in length (Figure 2).

**Table 1.** Summary statistics of *G. lacustris* gonad transcriptome sequencing data.

| Sample | Raw Reads | Clean reads | Clean bases | Error(%) | Q20(%) | Q30(%) | GC(%) |
| --- | --- | --- | --- | --- | --- | --- | --- |
| FG | 63,391,676 | 60,657,644 | 9.1G | 0.02 | 96.03 | 90.26 | 44.97 |
| MG | 54,542,800 | 52,016,136 | 7.8G | 0.02 | 96.54 | 91.05 | 47.95 |

Female *G. lacustris* named FG; male *G. lacustris* named MG.

**Figure 2.**
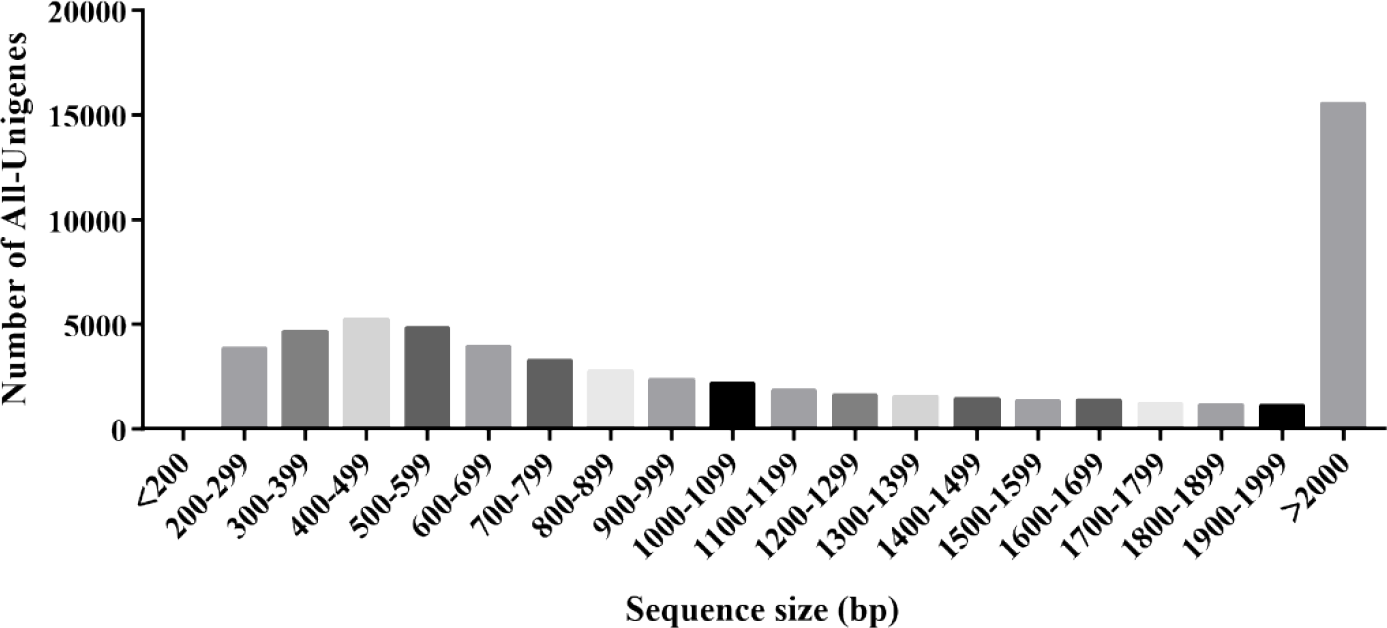
Length distribution of assembled unigenes

### 3.2. Sequence annotation and classification of the *G. lacustris* transcriptome

The 62,573 unigenes were searched against NR, NT, Swiss-Prot, COG/KOG, Pfam, GO and KEGG using BLAST (E value < 1E^−5^). Based on this search, 41,480 and 37,500 unigenes were classified in the NR and Swissprot databases, respectively, while 18,318 and 27,009 unigenes were assigned to the COG/KOG and KEGG databases, respectively. A total of 47,891 (76.53% of 62,573) unigenes had at least hit to one or more of the databases (Table s2). By searching against the NR database, we identified 40,726 unigenes that matched to known sequences from 236 species, including 26,379 unigenes that were homologous to *Boleophthalmus pectinirostris* (64.77%), 1,573 unigenes that were homologous to *Lates calcarifer* (3.77%) (Figure s1).

To better understand the functional categories of the genes involved in ovary and testis development and function in *G. lacustris*, the unigenes identified above were searched against GO, COG/KOG, and KEGG databases. In the GO analysis, 35,394 (56.56% of 62,573) unigenes were classified into three major GO categories including biological process, cellular component and molecular function (Figure 3). Among these functional groups, the terms “cellular process” (21,652, 61.17% of 35,394), “cell” (12,906, 36.46%), and “binding” (22,407, 63.31%) were dominant in the biological process, cellular component, and molecular function categories, respectively. In the COG/KOG database, 18,318 (29.27% of 62573) unigenes were annotated and classified into 26 functional categories (Figure 4). The largest functional category was signal transduction mechanisms (3,292, 17.97% of 18,318), followed by general function prediction only (3,035, 16.57%). According to the participating KEGG metabolic pathway, 27,009 (43.16% of 62,573) unigenes were divided into five branches: Cellular Processes, Environmental Information, Processing, Genetic Information Processing, Metabolism, and Organismal Systems. The major branches “Organismal Systems” consisted of 8,329 unigenes (30.84% of 27,009; Figure 5).

**Figure 3.**
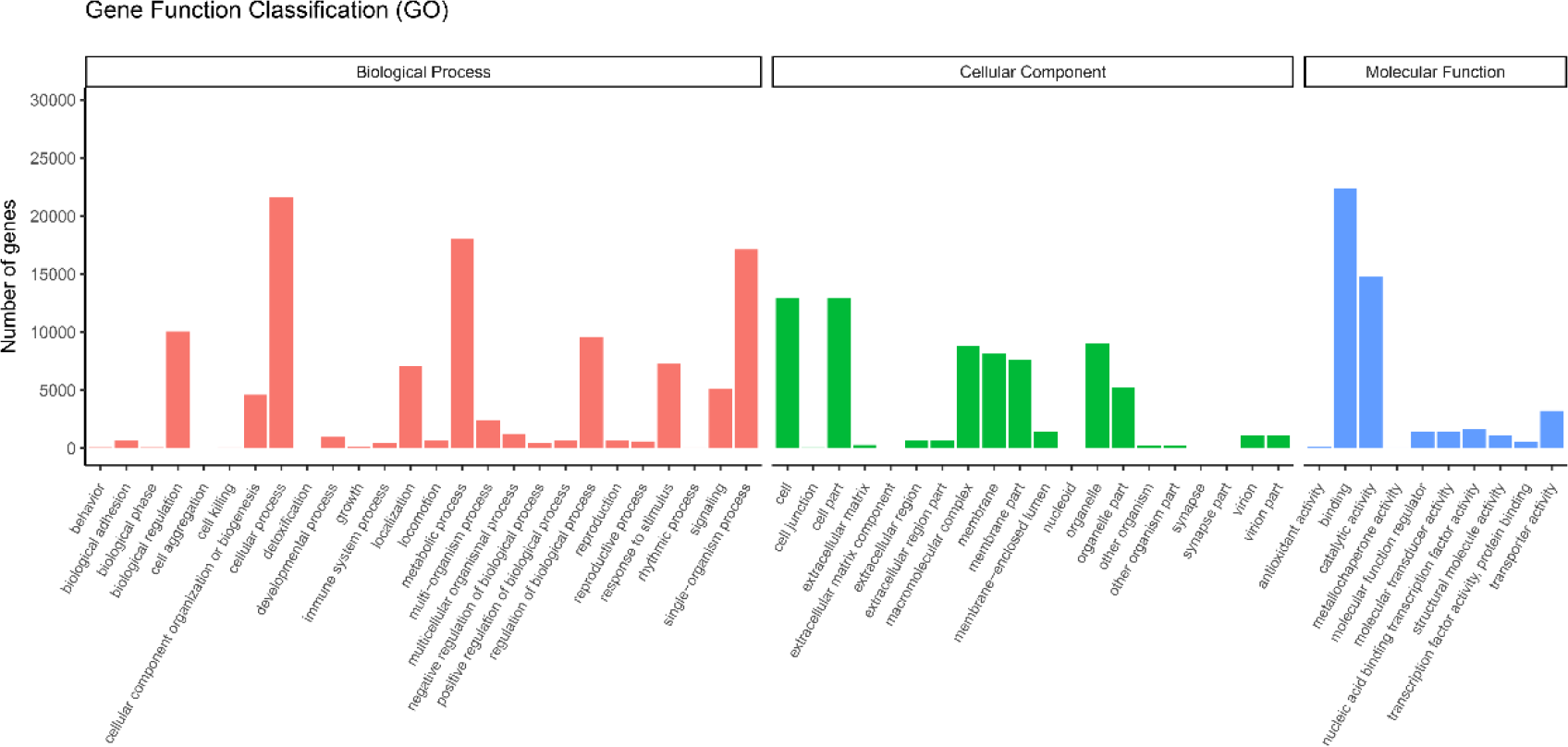
Gene ontology (GO) assignment of assembled unigenes of *G. lacustris*. The *x*-axis shows the GO functions and the *y*-axis shows the number of genes with the GO function.

**Figure 4.**
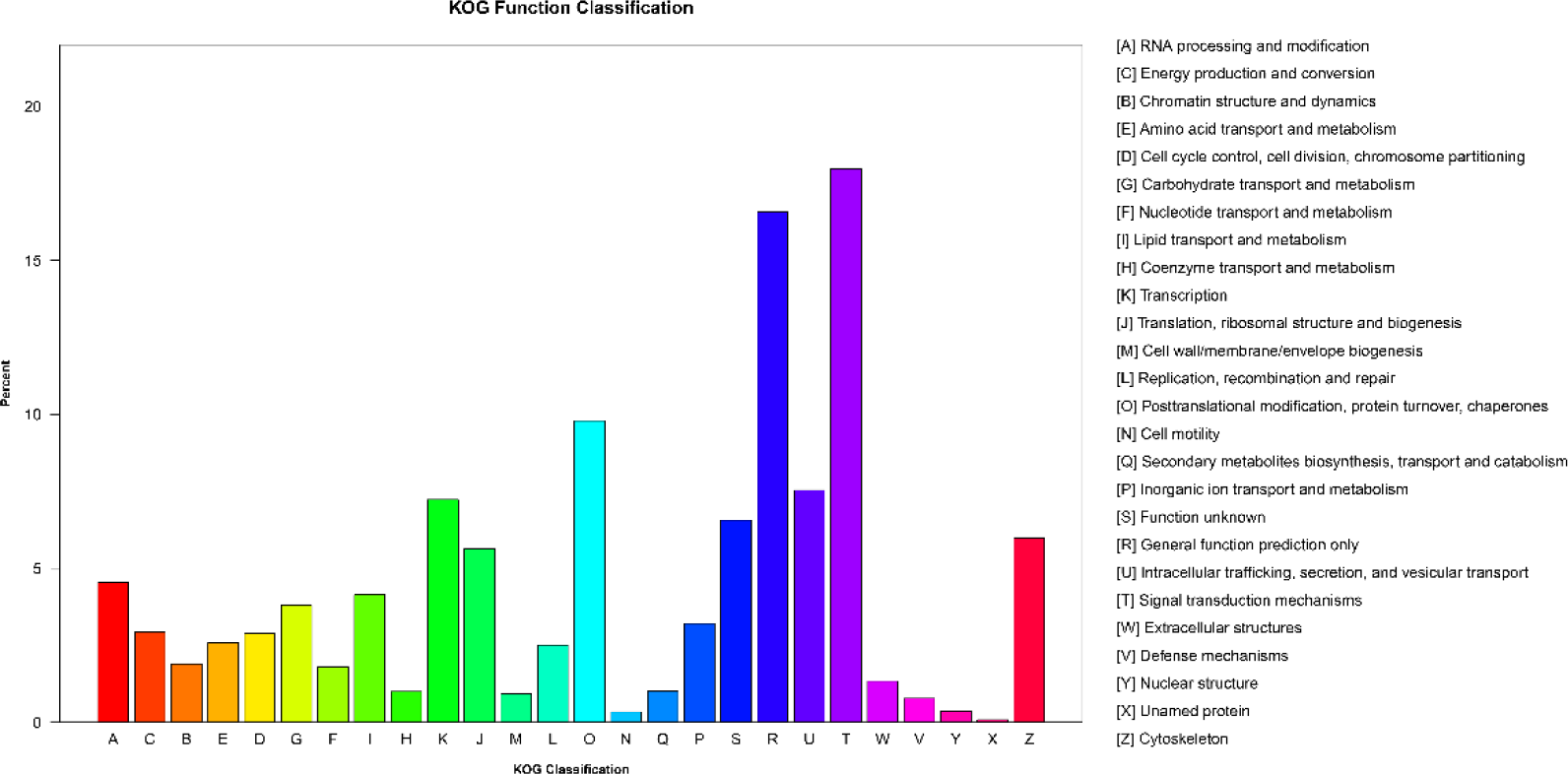
Clusters of Orthologous Groups (COG/KOG) function classification of unigenes in All-Unigene. *x*-axis, function classes of COG; *y*-axis, numbers of unigenes in one class. *Right*, the full names of the functions in *x*-axis.

**Figure 5.**
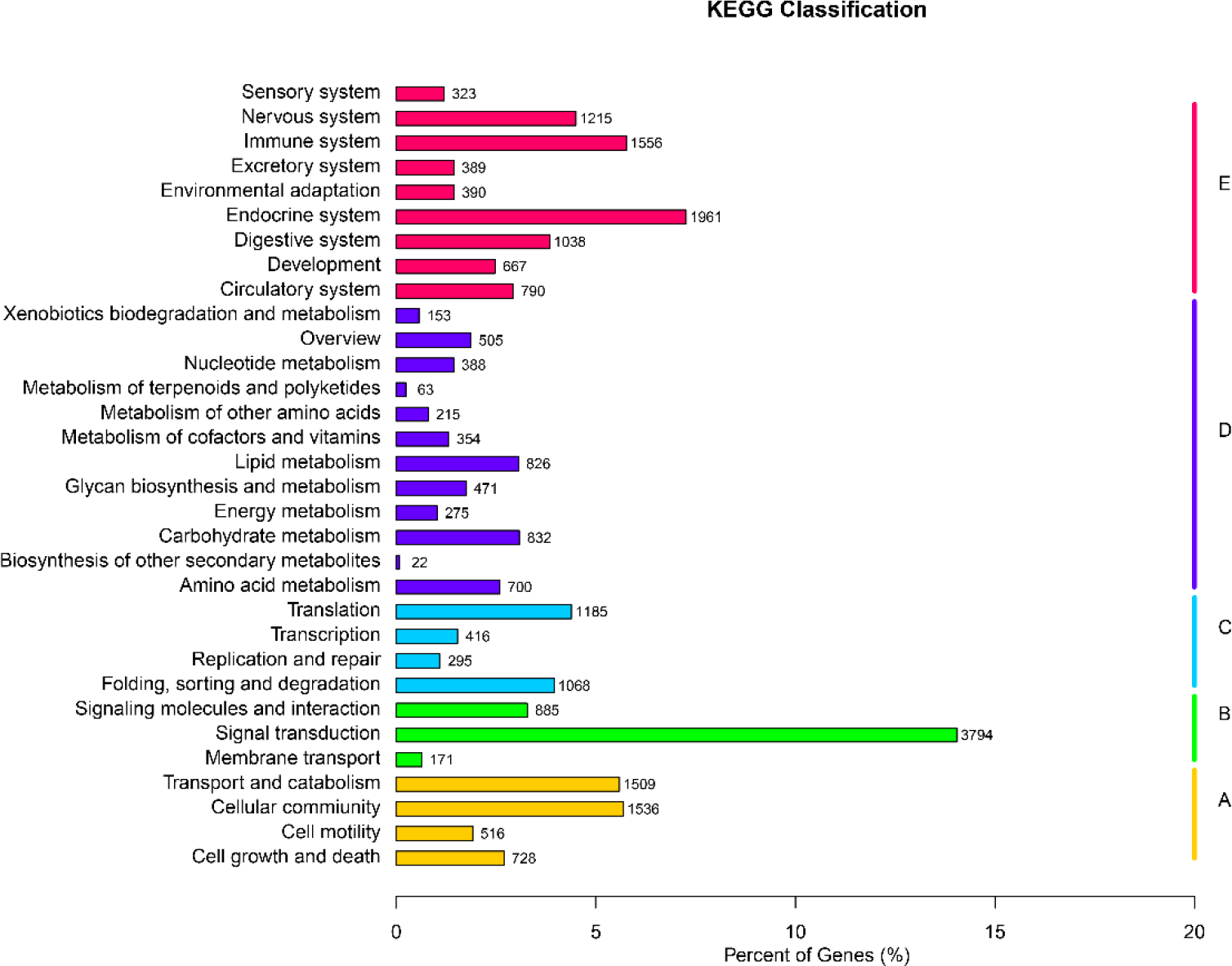
Description of the mapped Kyoto Encyclopedia of Genes and Genomes (KEGG) pathways. The *x*-axis labels the number of unigenes mapped into each KEGG pathway and the *y*-axis labels the distribution of the KEGG pathways. A, Cellular Processes; B, Environmental Information Processing; C, Genetic Information Processing; D, Metabolism; E, Organismal Systems.

### 3.3. Differential gene expression identification and enrichment analysis

Next, we identified genes that were enriched in the male and female gonads by comparing the male and female gonad transcriptomes using DEG analysis. Of the 56,735 unigenes expressed in the gonads, 32,643 unigenes were commonly expressed in the male and female, 13,628 were female-bised and 10,464 were male-biased (Figure s2). Meanwhile, 8,571 unigenes were up-regulated in the female and 2,383 unigenes were up-regulated in the male (FDR ≤ 0.005 and log_2_FC > 1) (scripts listed in Supplementary data s1). Many of the genes identified in the transcriptome were expressed in both male and females, and these DEGs included genes involved in sex determination, sex differentiation and gametogenesis, such as doublesex and mab-3 related transcription factor 1 (*dmrt1*), forkhead box protein L2 (*foxl2*), cytochrome P450 aromatase A (*cyp19a1a*), estrogen receptor alpha (*esra*), inhibin alpha, (*inha*)inhibin beta (*inhb*), testis development-related protein (*tdrp*) and GATA-type zinc finger protein 1 (*zglp1*).

The GO and KEGG analysis were employed to identify potential functions of DEGs. As shown in Figure 6, cellular metabolic process (4,199) was the most abundant GO function items in biological process, while cell (3,501) and cell part (3,501) were the most abundant in cellular component. In molecular function, binding (6,151) was enriched compared to others. Meanwhile, 3,618 DEGs mapped to 298 KEGG pathways including ubiquitin mediated proteolysis, ribosome biogenesis in eukaryotes, oocyte meiosis, progesterone-mediated oocyte maturation, TGF-beta signaling pathway, PI3K-Akt signaling pathway, Ovarian steroidogenesis, and steroid hormone biosynthesis (Supplementary data s2 & s3).

**Figure 6.**
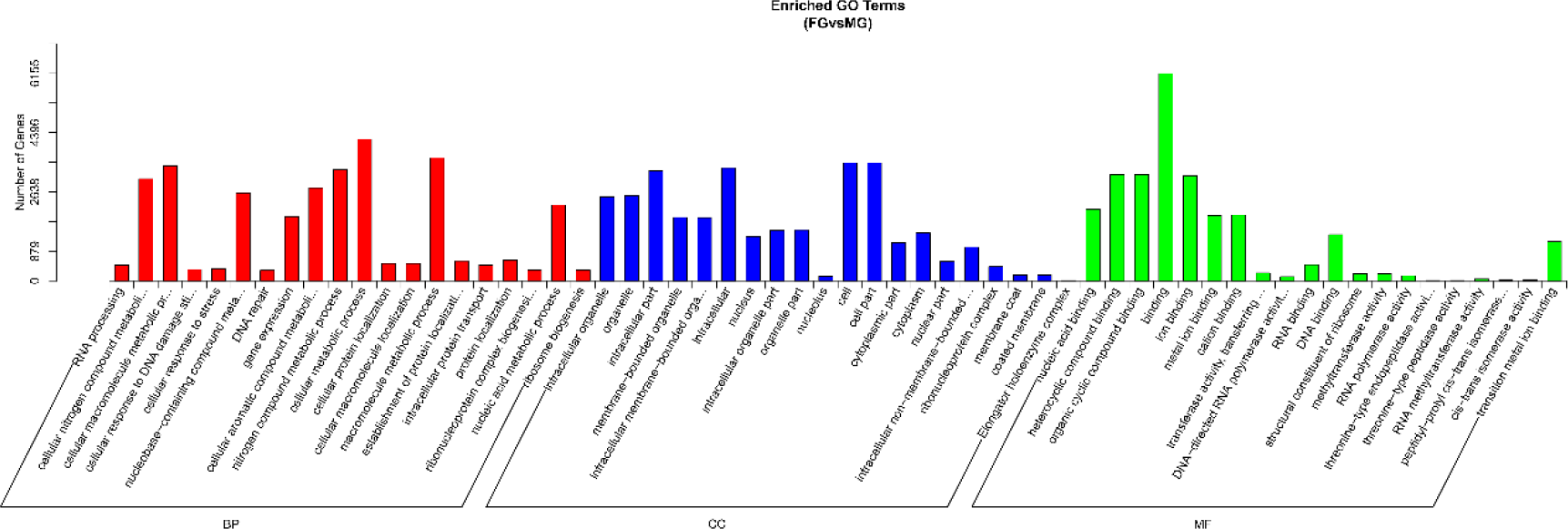
Gene ontology (GO) assignment of differentially expressed genes (DEGs) of *G. lacustris*. The *x*-axis labels GO functions and the *y*-axis labels the number of genes with GO function. FG: Female gonad; MG: Male gonad; BP: Biological Process; CC: Cellular Component; MF: Molecular Function

### 3.4. RT-qPCR validation

To validate the results of the transcriptomic analysis, 14 genes were selected for validation by RT-qPCR (Table 2). The expression patterns obtained using RT-qPCR analysis was consistent with the transcriptome analysis (Table 2).

**Table 2.** Validation of the RNA-seq data by RT-qPCR.

| Gene ID | Gene name | RNA-Seq |  | RT-qPCR |  |
| --- | --- | --- | --- | --- | --- |
|  |  | Log<sub>2</sub> (FG/MG) | q value (FG/MG) | Log<sub>2</sub> (FG/MG) | p value (FG/MG) |
| Cluster-9641.3059 | *cas3* | -6.69 | 4.88E-08 | -1.99 | 0.062 |
| Cluster-2179.0 | *cyp19a1a* | 3.75 | 0.012 | 1.75 | 0.016 |
| Cluster-9641.47528 | *dmrt1* | -7.12 | 2.25E-10 | -7.75 | 0.032 |
| Cluster-9641.47188 | *e3-1* | -8.81 | 2.77E-14 | -10.05 | 0.043 |
| Cluster-9641.16580 | *foxl2* | 4.73 | 0.00014 | 5.15 | 0.00018 |
| Cluster-9641.43183 | *gapdh* | -5.86 | 1.03E-72 | -4.5 | 0.022 |
| Cluster-9641.11433 | *inha* | 2.69 | 0.00070 | 1.5 | 0.023 |
| Cluster-9641.14038 | *inhb* | -4.95 | 0.00076 | -4.74 | 0.00052 |
| Cluster-9641.38934 | *krt2* | -7.48 | 5.37E-295 | -11.9 | 0.0058 |
| Cluster-9641.24002 | *rpl7* | -1.30 | 5.59E-24 | -0.72 | 0.20 |
| Cluster-9641.24530 | *sycp2* | 7.84 | 2.22E-20 | 6.9 | 0.0023 |
| Cluster-9641.6353 | *tdrp* | -3.4 | 0.30 | -2.22 | 0.067 |
| Cluster-9641.3821 | *test* | -5.39 | 6.19E-08 | -8.09 | 0.024 |
| Cluster-9641.25293 | *zglp1* | 9.61 | 1.89E-40 | 12.96 | 0.000035 |

Validation of RNA-Seq data using qRT-PCR. The RNA-Seq results are displayed as log2 (female read count /male read count). n = 3 for each sex. The results of qRT-PCR were performed by relative expression using *eef1b* and *hprt1* as the reference gene and measured by the method of optimized comparative Ct (2^−ΔΔCt^).

### 3.5. Characterization of simple sequence repeat (SSR) markers

Lastly, we used the MISA software to identify SSR markers for *G. lacustris* to genetic diversity research. In total, 38,550 SSRs were obtained from 20,517 (32.79 % of 62,573) unigenes (Table 3). Multiple SSRs were observed in 8,730 sequences, and 7,103 SSRs were present in compound formation. Six types of SSRs were detected, with the mono-nucleotide repeat motif comprising the largest group (17,419, 45.19%), followed by the di-nucleotide (13,121, 34.04%), tri-nucleotide (7,321, 18.99%), tetra-nucleotide (547, 1.42%), penta-nucleotide (87, 0.23%), and hexa-nucleotide (55, 0.14%) (Table 3). The relative abundance of different repeat motifs varied widely. A/T (16,671 of 17,419) was the most frequent motif in the mono-nucleotide SSRs. Among the di-nucleotide SSRs, AG/CT (8,591of 13,121) was most common, while GC/CG (2 of 13,121) was nearly undetectable. AGG/CCT (2,249 of 7,321), AAAT/ATTT (142 of 547), AAGGC/CCTTG (15 of 87) and ACCTGG/AGGTCC (19 of 55), were the most frequent motifs in tri-, tetra-, penta- and hexa- nucleotide SSRs, respectively. Specific primers can be designed according to the flanking sequence of these microsatellite loci.

**Table 3.** SSR summary from the *G. lacustris* transcriptome.

| Stat Item | numer |
| --- | --- |
| Total number of sequences examined | 62,573 |
| Total size of examined sequences (bp) | 116,923,694 |
| Total number of identified SSRs | 38,550 |
| Number of SSR containing sequences | 20,517 |
| Number of sequences containing more than 1 SSR | 8,730 |
| Number of SSRs present in compound formation | 7,103 |
| mono-nucleotide | 17,419 |
| di-nucleotide | 13,121 |
| tri-nucleotide | 7,321 |
| tetra-nucleotide | 547 |
| penta-nucleotide | 87 |
| hexa-nucleotide | 55 |

## 4. Discussion

Here, we report the first transcriptomic dataset for *G. lacustris*, focusing on the testis and ovarian tissues. To identify genes and biological pathways involved in gonadal development and reproduction, two cDNA libraries from ovaries and testes were prepared and sequenced on the Illumina platform. The N50 length is an important parameter for evaluating assembly quality, and the N50 length of our assembly (3082 bp) was higher than most published transcriptomes of aquatic animals, including *Paralichthys olivaceus* (809 bp) [9], *Patinopecten yessonsis* (565 bp) [31], *Larimichthys crocea* (1,934 bp) [32], *Oratosquilla oratoria* (1,502 bp) [33], *Andrias davidianus* (1,557 bp) [17] and *Sillago sihama* (2,190 bp) [14]. The annotation rate (76.53% of 62,573) of *G. lacustris* gonad transcriptomes was higher than the gonad transcriptomes of *C. feriatus* (37,500, 43.4% of 86,433) [34], *S. sihama* (34,104 of 74,038) [14] and *O. oratoria* (19,404 of 37,906) [33]. However, 14,682 (23.47% of 62,573) unigenes did not have any BLAST hits. This may be due to the limited genomic information available for *G. lacustris*, or because the unannotated unigenes contained 3’ or 5’ untranslated regions, non-coding RNAs, or some unigenes were new genes, or short sequences containing unknown protein domains [17, 35]. When searched against the NR database, 26,379 (64.77% of 40,726) unigenes were found to be homologous to *Boleophthalmus pectinirostris* (Figure s1), and this may be due to the close evolutionary relationship of *G. lacustris* and *B. pectinirostris*, both of which are gobies [36].

The GO categories “cellular process”, “cell” and “binding” were enriched in the gonadal transcriptome (Figure 3). Of the COG/KOG categories, most enriched functional category was signal transduction mechanisms (Figure 4). These functional categories are relatively similar to gonadal transcriptomes of aquatic animals, indicating that the genes that belong to these categories are highly conserved [13, 14, 17, 37, 38]. We also found male- and female-biased unigenes that were shared between *G. lacustris* and other species [13, 14, 31, 33], suggesting that the molecular mechanisms underlying gonadal differentiation is shared among these species.

### 4.1. Genes related to gonadal development and gametogenesis in *G. lacustris*

Of the DEGs that were identified between the male and female gonadal transcritpomes of *G. lacustris,* well established genes involved in gonadal development and gametogenesis were identified, including *dmrt1*, *foxl2*, *gtsf1, sycp2, soxs, zglp1*, *tdrp, cyp19a1a*, and *zps etc*.

Doublesex- and mab-3-related transcription factor 1 (*dmrt1*) is a sex determining gene that encodes a transcription factor that is essential for the maintenance of male-specified germ cells and testis differentiation [7, 39]. Forkhead box protein L2 (*foxl2*) is essential for ovarian differentiation and inhibits the male genetic pathway at several stages of female gonadal differentiation in vertebrates [40–43]. In *G. lacustris*, *dmrt1* is highly expressed in the testis, while *foxl2* is highly expressed in the ovary, and *foxl2* and *dmrt1* likely constitute an antagonistic regulation system during gonadal development. In mice, ectopic expression of *dmrt1* can account for almost all transcriptome changes caused by *foxl2* deletion [44]. In Nile tilapia, *foxl2* and *dmrt1* play antagonistic roles in sex differentiation by regulating *cyp19a1a* expression and estrogen production [45].

Sox (Sry-type HMG box) proteins regulate sex determination and differentiation in vertebrate. In mice, *sox5* and *sox6* is exclusively expressed in the testis and regulates spermatogenesis [46, 47], *sox9* regulate of testis organogenesis and germ cell maintenance [48, 49]. In contrast, found that only *sox9* is highly expressed in testis, while *sox5 sox6, sox10* and *sox19* were expressed at higher levels in the female gonads than in the male (Table s3), suggesting that the *sox* genes may play key roles in primarily ovary development in *G. lacustris*. In *Dicentrarchus labrax,* sox19 mRNA also was expressed at higher levels in the mature ovary compared to the mature testis [50].

In addition to *soxs*, we found high expression of *sycp2*, *zglp1*, *gtsf1, inha*, and *zps* in the ovary, while *inhb* and *tdrp* was highly expressed in the testis. Synaptonemal complex protein 2 (*sycp2*) is required for cohesin core integrity at diplotene during the meiotic division [51]. Lack of *sycp2* results in reduction of female fertility and spermatocyte apoptosis in mice [52]. GATA-type zinc finger protein 1 (*zglp1*) is a nuclear zinc finger protein that regulates the interaction between somatic cells and germ cells during gonad development in mammals [53, 54], while it may play an important role in oogenesis in *C. semilaevis* [55]. In mammals, gametocyte-specific factor 1-like (*gtsf1*) is highly expressed in embryonic male and female gonads, and is expressed in germ cells throughout development, suggesting that it may regulate germ cell development [56, 57]. In zebrafish, *gtsf1* plays a conserved role in vertebrate gametogenesis. The zona pellucida (*zp*) genes encode ZP glycoproteins that are involved in oocyte development and fertilization. On the zona pellucida (ZP), sperm receptor activity is associated with glycoproteins ZP3 and ZP2 [58]. In fish, *zps are* mainly expressed in the ovary, and we found higher expression levels for *zp2*, *zp3*, and *zp4* in the ovary (Table S3), suggesting that these *zps* may also play important roles in oocyte development and fertilization [14, 59, 60]. As a member of the TGF-β family, inhibins (*inha* and *inhb*) play important roles in the development of vertebrate, especially mammalian gametes [61]. Wu et al. (2000) found that *inha* regulates zebrafish oocyte maturation in a dose-dependent manner [62]. Lankford et al. (2010) found that rainbow trout *inha* is regulated by maturation-inducing hormone (MIH), and MIH regulates follicular maturation [63, 64]. Interestingly, we found high expression of *inha* found in the ovary and *inhb* in the testis, suggesting that inhibins may regulate gametogenesis in *G. lacustris*. In addition, testis development-related protein (*tdrp*) is a nuclear factor that is predominantly expressed in spermatogenic cells of the testis and regulates spermatogenesis [65]. These genes likely constitute a network that regulates gonadal development and gametogenesis in *G. lacustris*.

### 4.2. Candidate pathways in gonadal development and gametogenesis

We also identified different KEGG enrichment results between the testis and ovary of *G. lacustris*. Ubiquitin mediated proteolysis, cell cycle, oocyte meiosis, progesterone-mediated oocyte maturation, and p53 signaling pathway were significantly enriched in the ovary (Supplementary data s2), while PI3K-Akt signaling pathway, steroid hormone biosynthesis, drug metabolism - cytochrome P450 and metabolism of xenobiotics by cytochrome P450 were significantly enriched in the testis (Supplementary data s3). This suggests that different regulatory mechanisms exist between the ovary and testis of *G. lacustris*.

The development and maturation of the ovary is a complex process that involves several functional pathways [66]. We found that there were 117 putative proteins involved in the cell cycle, 104 involved in ubiquitin-mediated proteolysis, 75 involved in oocyte meiosis and 56 involved in progesterone-mediated oocyte maturation (Supplementary data s2). Similar findings were also found in *Portunus trituberculatus* [66, 67].

*p53* maintains integrity of biological processes and also regulate reproduction [13, 68]. In the zebrafish, *p53* regulates the number of germ cells and mediates sex-determination indirectly [69]. The p53 signaling pathway was significantly enriched in the ovary, suggesting that it may regulate *G. lacustris* reproduction [13].

Steroid hormones often determine the development of the fish reproductive system. Steroid hormones are primarily synthesized and secreted by the gonads and can be transported to other target tissues through the circulation of blood, and function in combination with nuclear receptors, such as estrogen receptors and androgen receptors [70]. The p450 family regulates steroid hormone biosynthesis. We found that steroid hormone biosynthesis, metabolism of xenobiotics by cytochrome P450 and drug metabolism - cytochrome P450 pathways were significantly enriched in the testis, suggesting that steroid-metabolizing enzymes may participate in the development of the testis and regulate the synthesis of steroid hormones.

The PI3K/AKT pathway is involved in testicular growth and development and can regulate the secretion of androgen and the production of sperm [71]. PI3K is an important class of lipid kinases that specifically catalyze phosphatidylinositol lipids and is the first regulator of AKT [72]. Activated PI3K transports AKT to the cell membrane to activate AKT. AKT is a serine/threonine protein kinase, and within the testis, AKT is primarily expressed in spermatogenic cells and supporting cells. Phosphorylated AKT promotes cell survival and proliferation and regulates apoptosis [73]. Here, PI3K-Akt signaling pathway was enriched in testis, suggesting that may regulate male reproductive development and maintain testicular homeostasis.

### 4.3. Discovery of SSRs

SSR is an effective method for analyzing DNA polymorphisms, which is mainly used in genetic mapping, marker-assisted plant breeding, and genome fingerprinting. With the continuous development of high-throughput sequencing technology, the isolation of EST-SSR markers using sequences obtained from transcriptome data has gradually become a low-cost and high-efficiency EST-SSR development technology. In the *G. lacustris* transcriptome, 38,550 SSRs were obtained from 62,573 unigenes. Among the SSRs that were identified, mono-nucleotide repeat motifs were the most abundant, and there were fewer SSRs with higher repeated nucleotides (Table 3), which is consistent with analysis of SSRs from spotted knifejaw, crucifix crab, Chinese giant salamander and silver sillago. AG/CT repeats were most common among the di-nucleotide SSRs (65.5%), and CG/GC repeats were the least common, accounting for 0.015% of all di-nucleotide repeats. Similar results were obtained in the study of the crucifix crab gonad transcriptome by Zhang et al. This may be due to the fact that the DNA double helix structure has two hydrogen bond phases between the AT bases, and three between CGs. The presence of CG/GC microsatellites may make the unraveling of the DNA double strands to be difficult and affect replication and transcription. These microsatellite markers will provide useful tools for investigating genetic population structure, population history and conservation management of *G. lacustris*.

## 5. Conclusions

Here, we report the first gonadal transcriptomes of *G. lacustris*. A total of 62,573 unigenes were obtained, and 47,891 unigenes were mapped to major databases, enriching the functional genomic resources for *G. lacustris*. By comparing ovary and testis transcriptomes, numerous genes (*foxl2*, *dmrt1*, *cyp19a1a*, *inha*, *inhb*, *sycp2*, *zglp1*, *zps*, *esra*, etc.) and pathways such as Ubiquitin mediated proteolysis, oocyte meiosis, progesterone-mediated oocyte maturation, p53 signaling pathway, PI3K-Akt signaling pathway and steroid hormone biosynthesis pathways that may be involved in gonadal development and gametogenesis genes were identified. In addition, 38,550 SSRs were obtained from 20,517 unigenes with SSR containing sequences. These findings lay the foundation for future functional analyses of *G. lacustris*.

## Supplementary Materials

Figure s1, Species classification by searching against the NR database.

Figure s2, FGvsMG.Expression_Venn_diagram.

Table s1, The sequences of primers in this study.

Table s2, Numbers of unigenes with annotation results in different databases.

Table s3, List of putative genes significantly expressed in male and female gonads of *G. lacustris*.

Supplementary data s1, FGvsMG.DEGannot.

Supplementary data s2, FGvsMG_up.DEG_KEGG_pathway_enrichment_result.

Supplementary data s3, FGvsMG_down.DEG_KEGG_pathway_enrichment_result.

## Author Contributions

**Conceptualization**: Zhongdian Dong, Chengqin Huang, and Zhongduo

**Data curation**: Zhongdian Dong, Chengqin Huang, Hairui Zhang, Shunkai Huang, Ning Zhang

**Formal analysis**: Zhongdian Dong, Changxu Tian, and Yusong Guo

**Funding acquisition**: Yusong Guo.

**Methodology**: Chengqin Huang, Hairui Zhang, Shunkai Huang, Ning Zhang

**Project administration**: Zhongdian Dong, Yusong Guo.

**Writing – original draft**: Zhongdian Dong.

